## Supplementary Information for "Identifying small-molecules binding sites in RNA conformational ensembles with SHAMAN"

#### Supplementary Tables

| system | PDB id holo | ligand id | PDB id apo | global RMSD [ Å ] | binding site RMSD [ Å ] |
|---|---|---|---|---|---|
| FMN riboswitch | 6dn3 | GZ4 | 6wjr | 1.82 | 1.09 |
| THF riboswitch | 4lvx | H4B, H4B | 7kd1 | 3.74 | 0.36, 2.00 |
| TPP riboswitch | 3d2v | PYI | N/A | - | - |
| dG riboswitch | 3ski | GNG | N/A | - | - |
| HIV-1 TAR | 1uts | P13 | 1anr | 5.59 | 3.82 |
| HCV-IRES-IIa | 3tzr | SS0 | 2nok | 7.43 | 3.97 |
| IAV promoter | 2lwk | 0EC | 1mfy | 4.40 | 0.20 |

**Table S1**. *SHAMAN benchmark set*. The first column lists the systems chosen for our benchmark, with the riboswitches and the viral RNAs highlighted in gold and violet, respectively. For each system, the columns with red and cyan headers report the PDB id of the experimentally resolved holo and (when available) apo structures, respectively, along with the ligand id in the holo structure. The last two columns report the RMSD between holo and apo structures calculated on the common RNA backbone atoms of the whole molecules and of the binding site region only. The THF riboswitch has two copies of the H4B ligand and therefore two values are reported for the binding site RMSD.

| starting PDB id | # nt | # water molecules | # structural ions | # extra ions [$K^+$/$CL^-$] | total # atoms | simulation time [μs * N] |
|---|---|---|---|---|---|---|
| 6dn3 | 109 | 17210 | 9 | 143/50 | 72552 | 1 * 19 |
| 6wjr | 111 | 15194 | 2 | 155/45 | 64554 | 1 * 19 |
| 4lvx | 89 | 12093 | 0 | 125/37 | 51405 | 1 * 17 |
| 7kd1 | 89 | 14005 | 2 | 133/41 | 59075 | 1 * 17 |
| 3d2v | 77 | 11284 | 1 | 97/33 | 47757 | 1 * 17 |
| 3ski | 67 | 8346 | 4 | 95/25 | 35663 | 1 * 17 |
| 1uts | 29 | 5423 | 0 | 44/16 | 22682 | 1 * 17 |
| 1anr | 29 | 5158 | 0 | 43/15 | 21620 | 1 * 17 |
| 3tzr | 36 | 6028 | 9 | 46/18 | 25346 | 1 * 17 |
| 2nok | 44 | 7988 | 7 | 51/23 | 33440 | 1 * 17 |
| 2lwk | 32 | 4862 | 0 | 45/14 | 20530 | 1 * 17 |
| 1mfy | 31 | 4515 | 0 | 43/13 | 19106 | 1 * 17 |

**Table S2**. *Details of the SHAMAN simulations.* For each of the 12 SHAMAN runs, we report the following details of the mother system simulations. From left to right: the PDB id of the starting structure (with the riboswitches and the viral RNAs indicated in gold and violet, respectively), the number of RNA nucleotides (nt), the number of water molecules, the number of structural ions present in the deposited PDB, the number of extra ions added to neutralize the simulation box at 0.15 M KCl concentration, the total number of atoms, the total simulation time (1 μs for mother and each replica system).

| FMN riboswitch | PDB id | # nt | ligand PDB id | buriedness [ au ] | deposition year | experimental technique | resolution [ Å ] | ref | HARIBOSS entry |
|---|---|---|---|---|---|---|---|---|---|
| pocket 1 | 2yie | 156 | FMN | 0.78 | 2011 | X-ray | 2.94 | DOI | 2yie |
| | 3f2q | 128 | FMN | 0.77 | 2008 | X-ray | 2.95 | DOI | 3f2q |
| | 3f2t | 130 | FMN | 0.77 | 2008 | X-ray | 3.00 | DOI | 3f2t |
| | 3f2w | 134 | FMN | 0.77 | 2008 | X-ray | 3.45 | DOI | 3f2w |
| | 3f2x | 129 | FMN | 0.77 | 2008 | X-ray | 3.11 | DOI | 3f2x |
| | 3f2y | 127 | FMN | 0.78 | 2008 | X-ray | 3.20 | DOI | 3f2y |
| | 3f4e | 127 | FMN | 0.77 | 2008 | X-ray | 3.05 | DOI | 3f4e |
| | 3f4g | 128 | RBF | 0.79 | 2008 | X-ray | 3.01 | DOI | 3f4g |
| | 3f4h | 125 | RS3 | 0.80 | 2008 | X-ray | 3.00 | DOI | 3f4h |
| | 3f30 | 128 | FMN | 0.79 | 2008 | X-ray | 3.15 | DOI | 3f30 |
| | 5c45 | 119 | 51B | 0.76 | 2015 | X-ray | 2.93 | DOI | 5c45 |
| | 5kx9 | 121 | 6YG | 0.75 | 2016 | X-ray | 2.90 | DOI | 5kx9 |
| | 6bfb | 124 | DKM | 0.77 | 2017 | X-ray | 2.82 | DOI | 6bfb |
| | 6dn1 | 147 | GZ7 | 0.76 | 2018 | X-ray | 3.03 | DOI | 6dn1 |
| | 6dn2 | 126 | GZG | 0.78 | 2018 | X-ray | 2.88 | DOI | 6dn2 |
| | 6dn3 | 124 | GZ4 | 0.76 | 2018 | X-ray | 2.80 | DOI | 6dn3 |

| THF riboswitch | PDB id | # nt | ligand PDB id | buriedness [ au ] | deposition year | experimental technique | resolution [ Å ] | ref | HARIBOSS entry |
|---|---|---|---|---|---|---|---|---|---|
| pocket 1, pocket 2 | 3sd3 | 90 | FFO | 0.62 | 2011 | X-ray | 1.95 | DOI | 3sd3 |
| | **4lvv** | **90** | **FFO** | **0.59, 0.57** | **2013** | **X-ray** | **2.10** | **DOI** | **4lvv** |
| | **4lvw** | **90** | **7DG** | **-** | **2013** | **X-ray** | **1.77** | **DOI** | **-** |
| | **4lvx** | **90** | **H4B** | **0.56, 0.60** | **2013** | **X-ray** | **1.90** | **DOI** | **4lvx** |
| | **4lvy** | **90** | **LYA** | **0.58, 0.64** | **2013** | **X-ray** | **2.00** | **DOI** | **4lvy** |
| | 4lvz | 90 | 6AP | - | 2013 | X-ray | 1.77 | DOI | - |
| | 4lw0 | 90 | ADE | - | 2014 | X-ray | 1.89 | DOI | - |

| TPP riboswitch | PDB id | # nt | ligand PDB id | buriedness [ au ] | deposition year | experimental technique | resolution [ Å ] | ref | HARIBOSS entry |
|---|---|---|---|---|---|---|---|---|---|
| pocket 1 | 2gdi | 79 | TPP | 0.76 | 2006 | X-ray | 2.05 | DOI | 2gdi |
| | 2hoj | 79 | TPP | 0.73 | 2006 | X-ray | 2.50 | DOI | 2hoj |
| | 2hok | 79 | TPP | - | 2006 | X-ray | 3.20 | DOI | - |
| | 2hol | 80 | TPP | 0.73 | 2006 | X-ray | 2.90 | DOI | 2hol |

| | | | | | | | | | |
|---|---|---|---|---|---|---|---|---|---|
| | 2hom | 81 | TPS | 0.75 | 2006 | X-ray | 2.89 | DOI | 2hom |
| | 2hoo | 84 | BFT | 0.75 | 2006 | X-ray | 3.00 | DOI | 2hoo |
| | 2hop | 77 | 218 | 0.70 | 2006 | X-ray | 3.30 | DOI | 2hop |
| | 3d2g | 78 | TPP | 0.74 | 2008 | X-ray | 2.25 | DOI | 3d2g |
| | 3d2v | 78 | PYI | 0.78 | 2008 | X-ray | 2.00 | DOI | 3d2v |
| | 3d2x | 78 | D2X | 0.75 | 2008 | X-ray | 2.50 | DOI | 3d2x |
| | 4nya | 79 | 2QB | 0.77 | 2013 | X-ray | 2.65 | DOI | 4nya |
| | 4nyb | 80 | 2QC | 0.75 | 2013 | X-ray | 3.10 | DOI | 4nyb |
| | 4nyc | 80 | SVN | - | 2014 | X-ray | 2.90 | DOI | - |
| | 4nyd | 80 | HPA | - | 2013 | X-ray | 3.15 | DOI | - |
| | 4nyg | 79 | VIB | 0.72 | 2013 | X-ray | 3.05 | DOI | 4nyg |
| dG riboswitch | PDB id | # nt | ligand PDB id | buriedness [ au ] | deposition year | experimental technique | resolution [ Å ] | ref | HARIBOSS entry |
| pocket 1 | 3ski | 68 | GNG | 0.81 | 2011 | X-ray | 2.30 | DOI | 3ski |
| | 3skl | 65 | GNG | 0.74 | 2011 | X-ray | 2.90 | DOI | 3skl |
| | 3skr | 65 | GNG | 0.78 | 2011 | X-ray | 3.10 | DOI | 3skr |
| | 3skt | 65 | GNG | 0.77 | 2011 | X-ray | 3.10 | DOI | 3skt |
| | 3skw | 65 | GNG | 0.75 | 2011 | X-ray | 2.95 | DOI | 3skw |
| | 3skz | 67 | GMP | 0.71 | 2011 | X-ray | 2.61 | DOI | 3skz |
| | 3slm | 67 | DGP | 0.73 | 2011 | X-ray | 2.70 | DOI | 3slm |
| | 3slq | 67 | 5GP | 0.70 | 2011 | X-ray | 2.50 | DOI | 3slq |
| pocket 2 | 3fo4 | 64 | 6GU | 0.77 | 2008 | X-ray | 1.90 | DOI | 3fo4 |
| | 3fo6 | 68 | 6GO | 0.79 | 2008 | X-ray | 1.90 | DOI | 3fo6 |
| | 3g4m | 68 | 2BP | - | 2009 | X-ray | 2.40 | DOI | - |
| | 3gao | 68 | XAN | - | 2010 | X-ray | 1.90 | DOI | - |
| | 3ger | 68 | 6GU | 0.81 | 2009 | X-ray | 1.70 | DOI | 3ger |
| | 3ges | 68 | 6GO | 0.79 | 2009 | X-ray | 2.15 | DOI | 3ges |
| | 3gog | 66 | 6GU | 0.76 | 2009 | X-ray | 2.10 | DOI | 3gog |
| | 3got | 68 | A2F | - | 2010 | X-ray | 1.95 | DOI | - |
| | 3rkf | 68 | DX4 | 0.83 | 2011 | X-ray | 2.50 | DOI | 3rkf |
| | 6ubu | 68 | GUN | - | 2019 | X-ray | 1.60 | DOI | - |
| | 6uc7 | 68 | Q44 | 0.69 | 2019 | X-ray | 1.80 | DOI | 6uc7 |
| | 6uc8 | 68 | ANG | 0.81 | 2019 | X-ray | 1.90 | DOI | 6uc8 |
| | 6uc9 | 68 | CMG | 0.73 | 2019 | X-ray | 1.94 | DOI | 6uc9 |

**Table S3**. *SHAMAN riboswitch validation set*. For each riboswitch in our benchmark set (gold cells), we report the details of the holo structures used for validation. Structures are grouped based on pocket and binding mode similarity (Materials and Methods). For each structure, we report from left to right: link to PDB entry, number of RNA nucleotides (nt), link to ligand PDB id, pocket buriedness (as calculated in HARIBOSS[1]), deposition year, experimental technique, resolution, link to reference publication, and link to HARIBOSS entry, if present. The THF riboswitch is sometimes resolved with two bound ligands (bold entries), therefore some properties are reported as a comma-separated list.

| HIV-1 TAR | PDB id | # nt | ligand PDB id | buriedness [ au ] | deposition year | experimental technique | resolution [ Å ] | ref | HARIBOSS entry |
|---|---|---|---|---|---|---|---|---|---|
| pocket 1 | 1uts | 30 | P13 | 0.55 | 2003 | NMR | - | DOI | 1uts |
| pocket 2 | 1arj | 30 | ARG | 0.67 | 1995 | NMR | - | DOI | 1arj |
| pocket 3 | 1lvj | 32 | PMZ | 0.61 | 2002 | NMR | - | DOI | 1lvj |
| pocket 4 | 1uud | 30 | P14 | 0.62 | 2003 | NMR | - | DOI | 1uud |
| | 1uui | 30 | P12 | - | 2003 | NMR | - | DOI | 1uui |
| pocket 5 | 2l8h | 30 | MV2003 | - | 2011 | NMR | - | DOI | - |
| HCV-IRES-IIa | PDB id | # nt | ligand PDB id | buriedness [ au ] | deposition year | experimental technique | resolution [ Å ] | ref | HARIBOSS entry |
| pocket 1 | 3tzr | 36 | SS0 | 0.66 | 2011 | X-ray | 2.21 | DOI | 3tzr |
| pocket 2 | 2ku0 | 39 | ISI | 0.57 | 2010 | NMR | - | DOI | 2ku0 |
| | 2ktz | 39 | ISH | 0.59 | 2010 | NMR | - | DOI | 2ktz |
| IAV promoter | PDB id | # nt | ligand PDB id | buriedness [ au ] | deposition year | experimental technique | resolution [ Å ] | ref | HARIBOSS entry |
| pocket 1 | 2lwk | 33 | 0EC | - | 2012 | NMR | - | DOI | 2lwk |

**Table S4**. *SHAMAN viral RNAs validation set*. Columns as defined in Tab. S3.

| FMN riboswitch | ΔΔG [ kJ/mol ] | rank | best match | distance [ Å ] | ΔΔG [ kJ/mol ] | rank | best match | distance [ Å ] |
|---|---|---|---|---|---|---|---|---|
| pocket 1 | < 0.1 | 2 | 1-1 04-DMEE 5kx9 6YG | 1.49 | < 0.1 | 2 | 0-1 05-MEPY 6dn3 GZ4 | 1.73 |
| THF riboswitch | ΔΔG [ kJ/mol ] | rank | best match | distance [ Å ] | ΔΔG [ kJ/mol ] | rank | best match | distance [ Å ] |
| pocket 1 | < 0.1 | 2 | 1-1 04-DMEE 5kx9 6YG | 2.37 | < 0.1 | 2 | 0-1 05-MEPY 6dn3 GZ4 | 1.51 |
| pocket 2 | < 0.2 | 3 | 1-1 04-DMEE 5kx9 6YG | 1.48 | < 0.2 | 3 | 0-1 05-MEPY 6dn3 GZ5 | 2.31 |
| TPP riboswitch | ΔΔG [ kJ/mol ] | rank | best match | distance [ Å ] | ΔΔG [ kJ/mol ] | rank | best match | distance [ Å ] |
| pocket 1 | 0.0 | 1 | 0-1 05-MEPY 4nyd HPA | 0.65 | - | - | - | - |
| dG riboswitch | ΔΔG [ kJ/mol ] | rank | best match | distance [ Å ] | ΔΔG [ kJ/mol ] | rank | best match | distance [ Å ] |
| pocket 1 | < 0.1 | 4 | 0-2 06-BENF 3slq 5GP | 1.57 | - | - | - | - |
| pocket 2 | 3.3 | 41 | 2-3 01-BENX 5kx9 6YG | 3.27 | - | - | - | - |

**Table S5.** *Details of the SHAMAPS corresponding to the experimental binding sites in the riboswitch benchmark set.* For each system and unique pocket, from left to right: ΔΔG (Eq. 9) of the SHAMAP with best overlap with the ligand in the experimental structures (Materials and Methods), rank, details of the interacting site with the best match (interacting site id with index of the RNA cluster, probe name, PDB id of the matching experimental structure, ligand name), and validation distance (Eq. 10). Columns with red and cyan headers report the results of the SHAMAN simulations initiated from holo-like and (when available) apo structures, respectively.

| HIV-1 TAR RNA | ΔΔG [ kJ/mol ] | rank | best match | distance [ Å ] | ΔΔG [ kJ/mol ] | rank | best match | distance [ Å ] |
|---|---|---|---|---|---|---|---|---|
| pocket 1 | 5.0 | 47 | 8-2 10-FORM 1arj ARG | 4.6 | 1.0 | 23 | 2-4 09-PIRZ 1arj ARG | 3.89 |
| pocket 2 | 0.2 | 11 | 0-8 08-IMIA 1lvj PMZ | 2.4 | 0.2 | 8 | 0-3 04-DMEE 1lvj PMZ | 2.94 |
| pocket 3 | 1.9 | 27 | 2-8 16-FORM 1uts P13 | 2.32 | 0.2 | 8 | 0-3 04-DMEE 1uts P13 | 6.25 |
| pocket 4 | 5.0 | 47 | 8-2 10-FORM 1uui P12 | 5.38 | 1.0 | 23 | 2-4 09-PIRZ 1uui P12 | 3.84 |
| pocket 5 | 5.0 | 47 | 8-2 10-FORM 2l8h MV2003 | 5.45 | 0.1 | 5 | 0-1 12-MAMY 2l8h MV2003 | 3.76 |

| HCV-IRES IIa | ΔΔG [ kJ/mol ] | rank | best match | distance [ Å ] | ΔΔG [ kJ/mol ] | rank | best match | distance [ Å ] |
|---|---|---|---|---|---|---|---|---|
| pocket 1 | 0.2 | 3 | 0-3 13-ACEY 3tzr SS0 | 2.67 | 2.9 | 14 | 1-1 14-MAMY-ions 3tzr SS0 | 2.84 |
| pocket 2 | 0.0 | 1 | 0-1 02-BETH 2ku0 ISI | 2.31 | < 0.1 | 5 | 0-1 08-IMIA 2ktz ISH | 3.17 |

| IAV RNA promoter | ΔΔG [ kJ/mol ] | rank | best match | distance [ Å ] | ΔΔG [ kJ/mol ] | rank | best match | distance [ Å ] |
|---|---|---|---|---|---|---|---|---|
| pocket 1 | 0.2 | 3 | 0-1 15-IMIA 2lwk 0EC | 1.43 | 5.9 | 47 | 1-3 13-ACEY 2lwk 0EC | 3.64 |

**Table S6.** *Details of the SHAMAPS corresponding to the experimental binding sites in the viral RNA benchmark set.* Columns as in Tab. S5.

| Probe id | 2D structure | Full name | Color |
|---|---|---|---|
| ACEY | 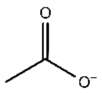 | Acetate | |
| BENX | 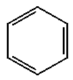 | Benzene | |
| DMEE | 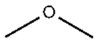 | Dymethil ether | |
| FORM | 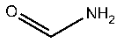 | Formamide | |
| IMIA | 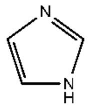 | Imidazole | |
| MAMY | 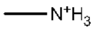 | Methyl-<br>ammonium | |
| MEOH | 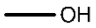 | Methanol | |
| PRPX | 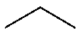 | Propane | |

**Table S7**. *First set of SHAMAN probes*. These probes were used in the development of SILCS-RNA[2]. From left to right: probe id, 2D structure, probe name, and color code.

| **Probe id** | **2D structure** | **Full name** | **Color** |
|---|---|---|---|
| BENX | 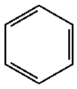 | Benzene | |
| BENF | 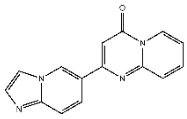 | dihydro-pyrido-pyrimidinone-imidazo-pyridine | |
| BETH | 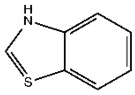 | Benzothiophene | |
| MEPY | 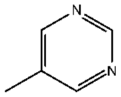 | Methyil-pyrimidine | |
| PIRZ | 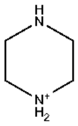 | Piperazine | |
| PYRD | 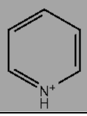 | Pyrimidine | - |

**Table S8**. *Second set of SHAMAN probes*. Probes determined in this work from the fragmentation of known RNA binders (Materials and Methods). Except for PYRD (dark grey), these constituted the second set of probes used in our SHAMAN simulations. From left to right: probe id, 2D structure, probe name, and color code.

| entire ligand | $P_{CC}$ | p-value | | Murcko scaffold | $P_{CC}$ | p-value |
|---|---|---|---|---|---|---|
| molecular weight | -0.08 | 0.70 | | molecular weight | 0.16 | 0.40 |
| # aromatic rings | 0.20 | 0.31 | | # aromatic rings | 0.19 | 0.32 |
| # H-bond donors | -0.21 | 0.28 | | # H-bond donors | 0.14 | 0.45 |
| # H-bond acceptors | -0.16 | 0.24 | | # H-bond acceptors | 0.05 | 0.82 |
| topological polar surface area | -0.24 | 0.21 | | topological polar surface area | 0.76 | 0.70 |
| # heterocycles | 0.34 | 0.08 | | # heterocycles | 0.36 | 0.08 |

**Table S9**. *Correlation between physico-chemical properties of ligands and successful probes.* For each property: name of the property, Pearson correlation coefficient ($P_{CC}$) and the corresponding p-value calculated between ligands and probes that successfully identified the corresponding binding site. Only SHAMAN simulations initiated from holo-like structures were considered for this analysis. Values were calculated considering either the entire ligand (left) or its Murcko scaffold (right).

| | **successful probes** | **unsuccessful probes** |
|---|---|---|
| **similar probes** | TP = 4 | FN = 9 |
| **dissimilar probes** | FP = 18 | TN = 42 |

| **Total population** | 73 |
|---|---|
| **TPR** | 0.31 |
| **TNR** | 0.70 |
| **PPV** | 0.08 |
| **NPV** | 0.82 |

**Table S10**. *Analysis of the relation between probe-ligand similarity and being a successful probe.* (Left) Confusion matrix to test the hypothesis that probes similar to ligands are successful, with number of cases with similar/dissimilar probes and successful/unsuccessful probes. Only SHAMAN simulations initiated from holo-like structures were considered for this analysis. (Right) True positive rate (TPR), true negative rate (TNR), positive predictive value (PPV), and negative predictive value (NPV).

| | 2yie | 3f2q | 3f2t | 3f2w | 3f2x | 3f2y | 3f4e | 3f4g | 3f4h | 3f30 | 5c45 | 5kx9 | 6bfb | 6dn1 | 6dn2 | 6dn3 |
|---|---|---|---|---|---|---|---|---|---|---|---|---|---|---|---|---|
| **2yie** | | 0.55 | 0.58 | 0.67 | 0.57 | 0.57 | 0.41 | 0.61 | 0.76 | 0.60 | 0.75 | 0.75 | 0.74 | 0.31 | 0.65 | 0.81 |
| **3f2q** | | | 0.48 | 0.53 | 0.35 | 0.32 | 0.36 | 0.56 | 0.66 | 0.54 | 0.66 | 0.70 | 0.60 | 0.69 | 0.55 | 0.70 |
| **3f2t** | | | | 0.47 | 0.44 | 0.50 | 0.49 | 0.61 | 0.64 | 0.43 | 0.69 | 0.67 | 0.74 | 0.57 | 0.56 | 0.65 |
| **3f2w** | | | | | 0.50 | 0.50 | 0.50 | 0.69 | 0.75 | 0.55 | 0.73 | 0.70 | 0.76 | 0.66 | 0.65 | 0.68 |
| **3f2x** | | | | | | 0.34 | 0.41 | 0.55 | 0.64 | 0.57 | 0.68 | 0.59 | 0.69 | 0.58 | 0.50 | 0.64 |
| **3f2y** | | | | | | | 0.44 | 0.58 | 0.63 | 0.55 | 0.73 | 0.64 | 0.66 | 0.63 | 0.57 | 0.77 |
| **3f4e** | | | | | | | | 0.52 | 0.63 | 0.46 | 0.71 | 0.61 | 0.73 | 0.50 | 0.51 | 0.69 |
| **3f4g** | | | | | | | | | 0.31 | 0.62 | 0.69 | 0.51 | 0.56 | 0.67 | 0.58 | 0.77 |
| **3f4h** | | | | | | | | | | 0.65 | 0.70 | 0.58 | 0.51 | 0.72 | 0.66 | 0.85 |
| **3f30** | | | | | | | | | | | 0.79 | 0.73 | 0.70 | 0.60 | 0.59 | 0.73 |
| **5c45** | | | | | | | | | | | | 0.39 | 0.69 | 0.78 | 0.69 | 0.66 |
| **5kx9** | | | | | | | | | | | | | 0.55 | 0.79 | 0.63 | 0.56 |
| **6bfb** | | | | | | | | | | | | | | 0.68 | 0.60 | 0.74 |
| **6dn1** | | | | | | | | | | | | | | | 0.62 | 0.74 |
| **6dn2** | | | | | | | | | | | | | | | | 0.59 |
| **6dn3** | | | | | | | | | | | | | | | | |

**Table S11**. *Similarity between the experimentally determined structures of the FMN riboswitch.* RMSD between all pairs of FMN riboswitch structures present in our validation set (Tab. S3). The RMSD was calculated on all the matching pairs of atoms of each pair of RNA molecules. All values are in Angstrom.

#### Supplementary Figures

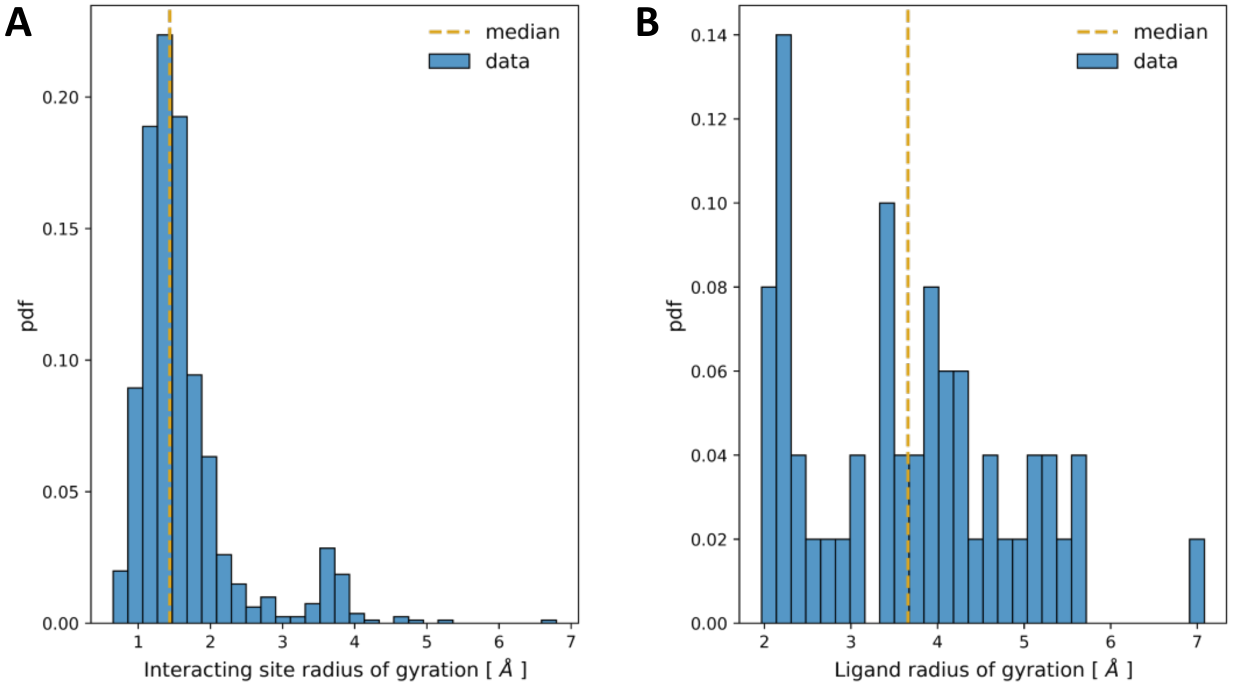

**Figure S1**. *Radius of gyration of SHAMAN interacting sites and ligands.* **A)** Normalized distribution of the radius of gyration of all the interacting sites detected by SHAMAN in all the systems of our benchmark set (Tab. S1). **B)** Normalized distribution of the radius of gyration of all the unique ligands present in the structures of our validation set (Tab. S3 and S4). In both plots, the median of the distribution is reported as a yellow dashed line.

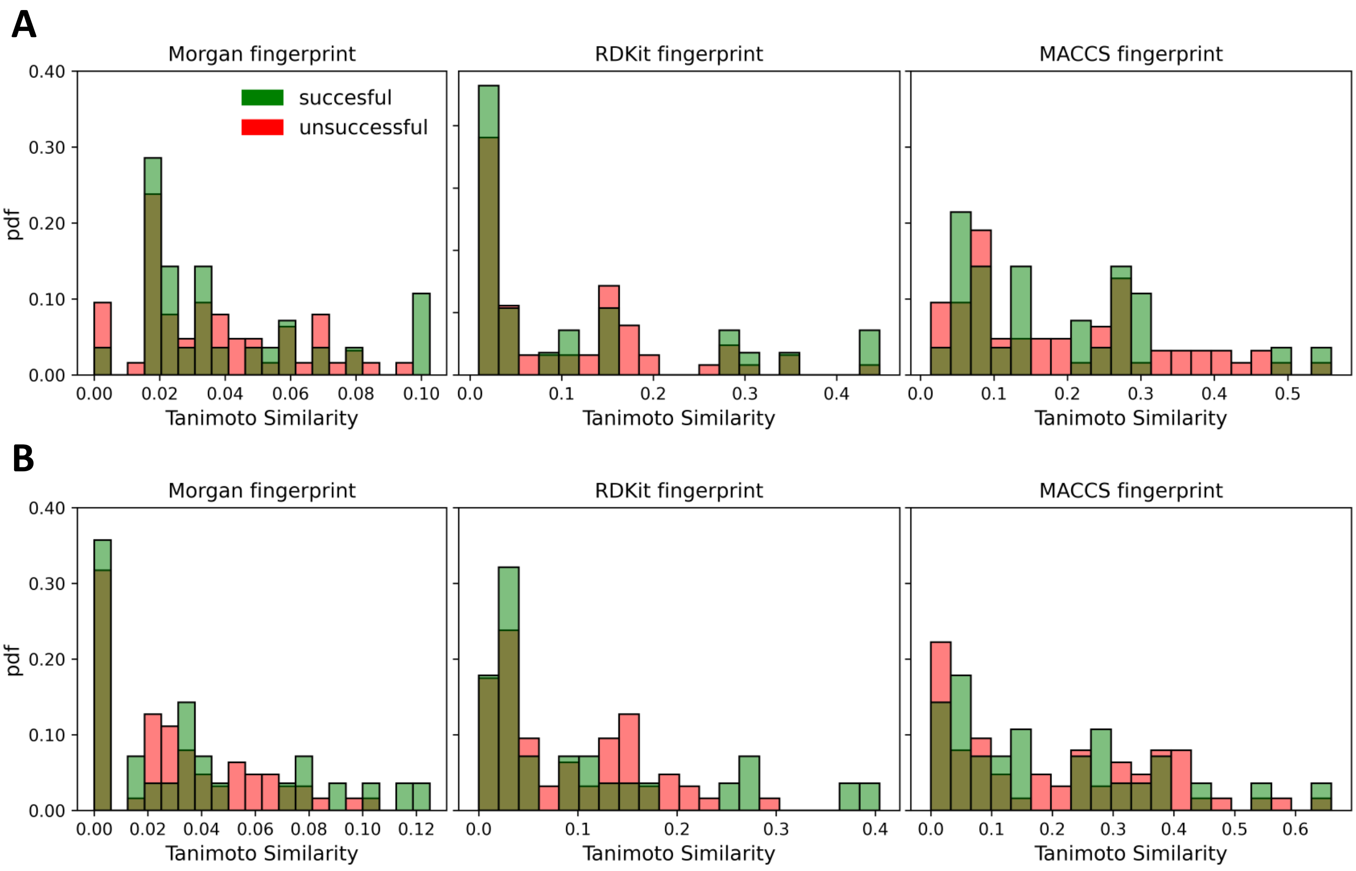

**Figure S2**. *Analysis of the similarity between ligands and probes.* **A)** Distributions of the Tanimoto similarity between the ligands present in the experimental structures of our benchmark set (Tab. S1) and the probes. Successful probes that identified the ligand are colored in green, unsuccessful probes in red. The analysis is limited to the SHAMAN simulations initiated from holo-like structures. From left to right, the analysis is performed with the 3 different fingerprints: Morgan, RDKit, MACCS. **B)** As in panel A, with similarities calculated using the Murcko scaffold instead of the entire ligand.

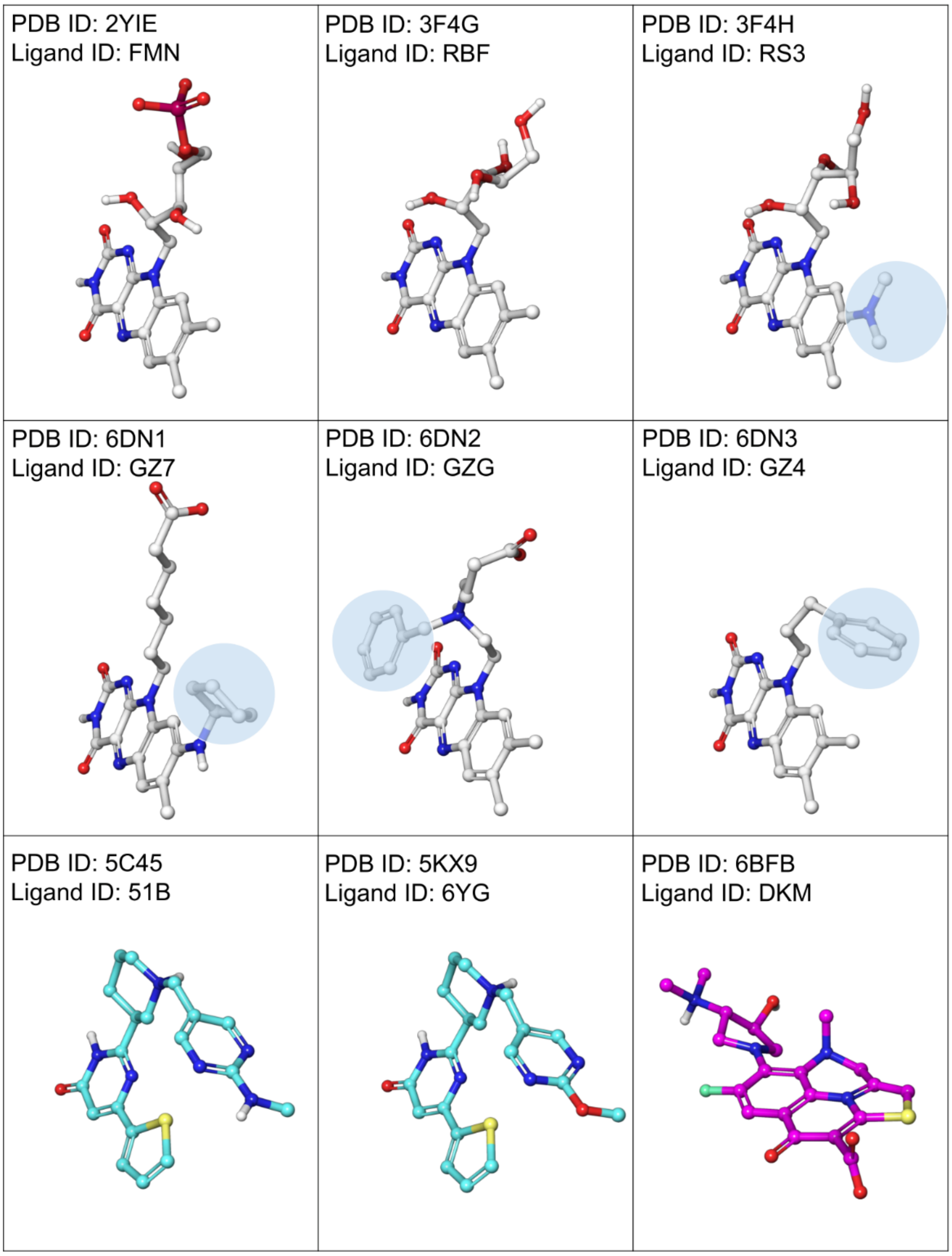

**Figure S3**. *Conformers of the FMN riboswitch binders.* The 3D conformers of the 9 unique ligands of the validation set of the FMN riboswitch are shown in each panel (Tab. S3). Following previous studies[3], we subdivided the FMN binders into 3 chemical families: the FMN, ribocil, and DKM families. The carbon atoms of the 3 families are reported in light grey, cyan and violet, respectively. The remaining atoms are reported with standard CPK colors. Significantly different chemical decorations among the members of the FMN family are highlighted by cyan circles. All ligands are aligned to facilitate the comparison between their 3D conformations. Only polar hydrogens are shown for clarity.

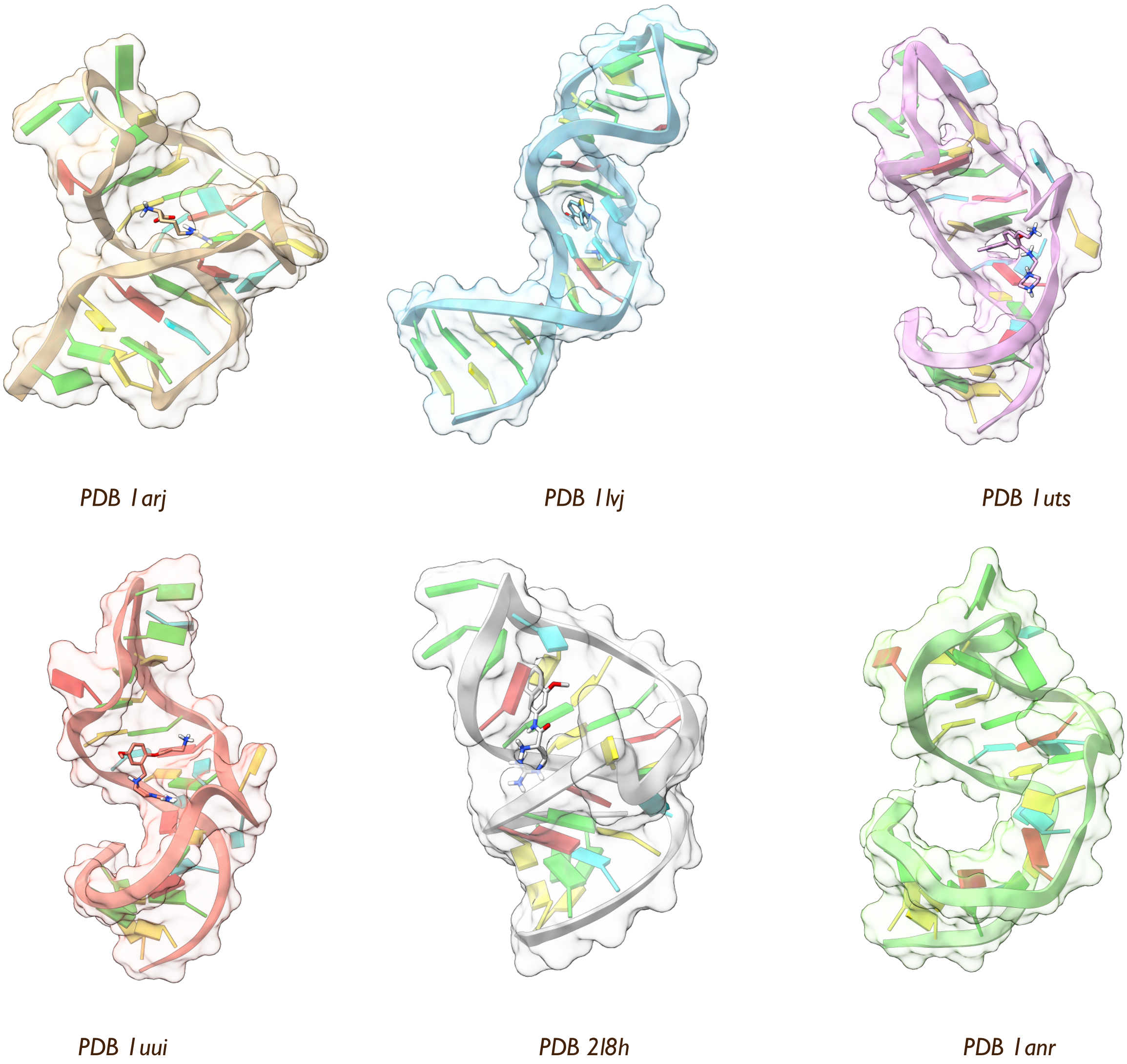

**Figure S4**. *Structural diversity of the HIV-1 TAR RNA.* Surface-ribbon representation of the structures of the 6 different conformations of the HIV-1 TAR RNA analyzed in this work along with the corresponding PDB id. From left to right and from top to bottom, the first 5 panels report the holo structures used in our validation set (Tab. S4) with the corresponding ligand highlighted in ball-and-stick representation. In the last panel, we illustrate an apo conformation of HIV-1 TAR.

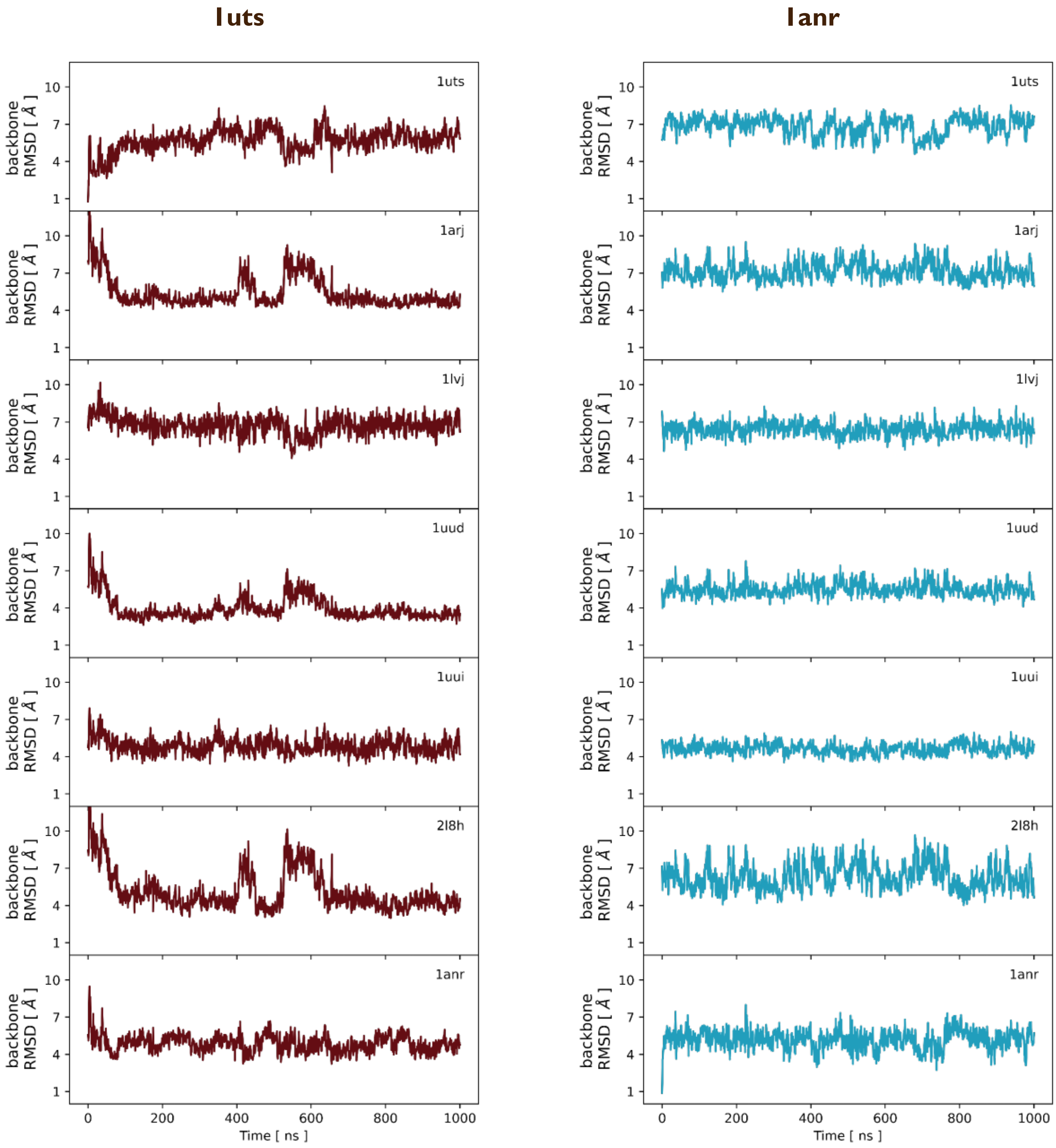

**Figure S5**. *Structural variety of the HIV-1 TAR RNA ensembles explored by SHAMAN.* RMSD of the backbone atoms of the HIV-1 TAR RNA in the SHAMAN mother simulation with respect to various experimental structures present in our validation set (Tab. S4), as a function of simulation time. Each panel corresponds to a different experimental structure, whose PDB id is indicated in the upper right corner. Simulations initiated from holo-like (PDB 1uts) and apo (PDB 1anr) conformations are highlighted in brown (left) and cyan (right), respectively.

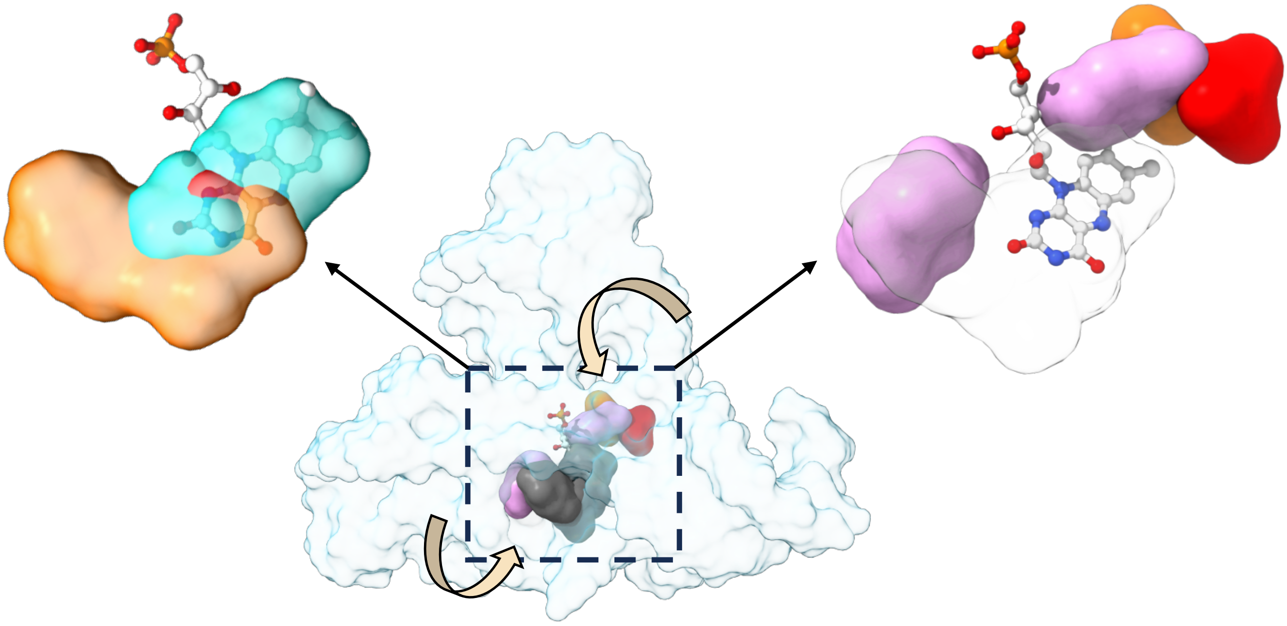

**Figure S6**. *Potential applications of SHAMAN in CADD*. In the center, the molecular surface of the most populated conformation of the FMN riboswitch obtained in the SHAMAN simulation initiated from an apo structure (PDB 6wjr). The dashed box indicates the FMN resolved binding site. The FMN ligand (PDB 2yie) as well as the SHAMAP that identified this binding site are superimposed by aligning the coordinates to the RNA cluster center. Different colors indicate different probes (color code as in Tab. S7 and S8). In the left panel, probe interacting sites of which the SHAMAP is composed of and their overlap with the ligand. In the right panel, interacting sites adjacent to the ligand binding site, with arrows suggesting two possible pathways to access the binding site.

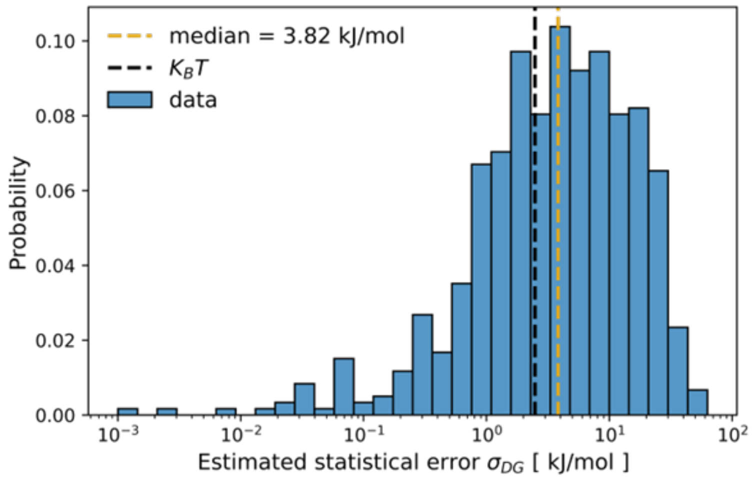

**Figure S7**. *Statistical error in the free energy of the probes interacting sites.* The normalized distribution of the calculated statistical error $\sigma_{\Delta G}^{l}$ on the free energy of each probe interacting site $l$ (Materials and Methods) calculated across all probes and systems. The median value of this distribution (3.82 $kJ/mol$), is reported as a yellow dashed line.

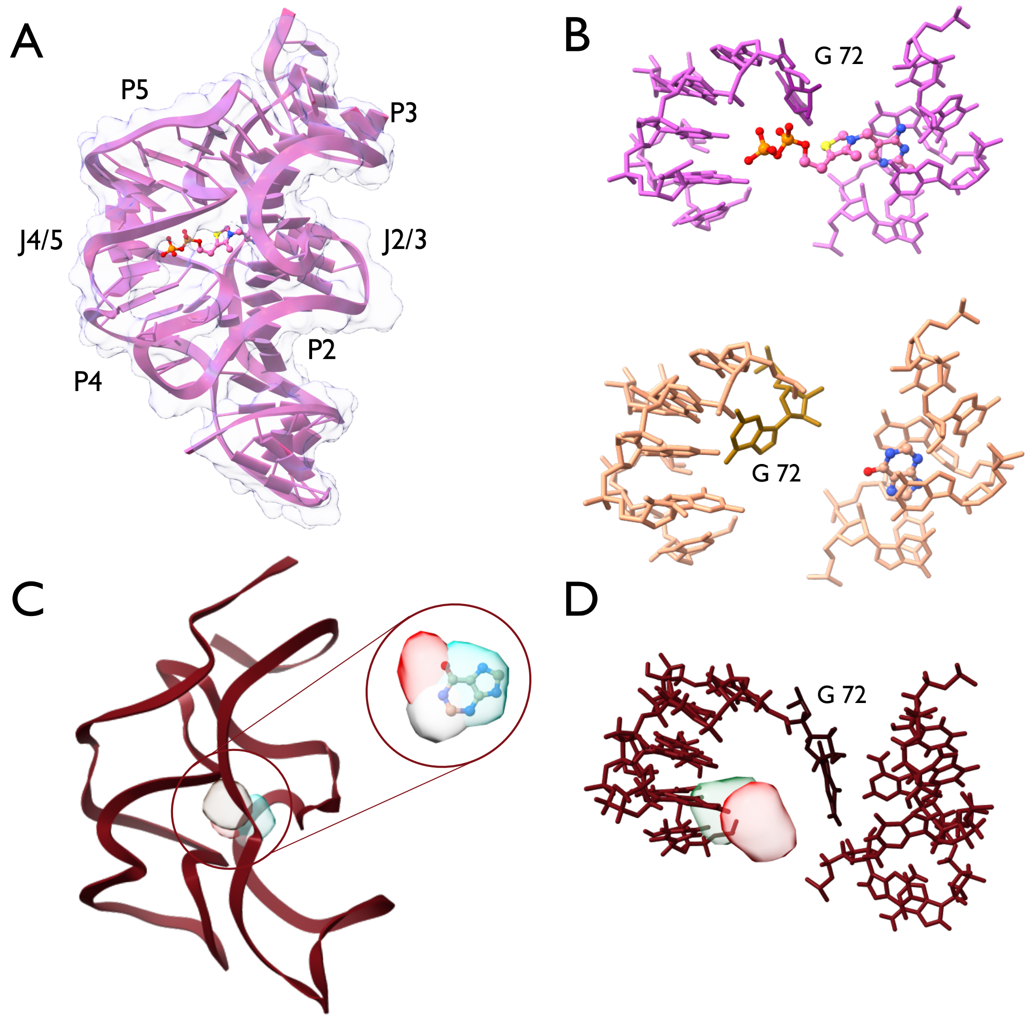

**Figure S8.** *The case of the TPP riboswitch.* **A)** Cartoon-surface representation of the TPP riboswitch in the bound conformation, as in PDB 2hoj[4]. P2-P5 indicate the helical regions, while J2/3 and J4/5 the multi-way junctions. **B)** In the upper panel, the TPP binding mode, as in PDB 2hoj[4]. In the lower panel, the binding mode of HPA, one of the fragments screened in the work related to PDB 4nyd[5]. **C)** On the left, a cartoon representation of the main conformation explored during SHAMAN simulations started from the holo-like state of the TPP riboswitch. The SHAMAPs identifying the experimental binding site are visualized as solid surfaces with the color code defined in Tab. S7 and S8. In the inset, the superposition of the mentioned SHAMAPs with ligand HPA, represented in sticks with CPK standard colors. Hydrogen atoms are visualized only when resolved in the experimental structure. **D)** The position of the interacting residues defined in **B** in the main conformation of TPP riboswitch explored during SHAMAN simulations. In panel **B** and **D**, the key residue G72 is highlighted by a darker color. The RNA nucleotides and ligands are represented using licorice and CPK styles, respectively.

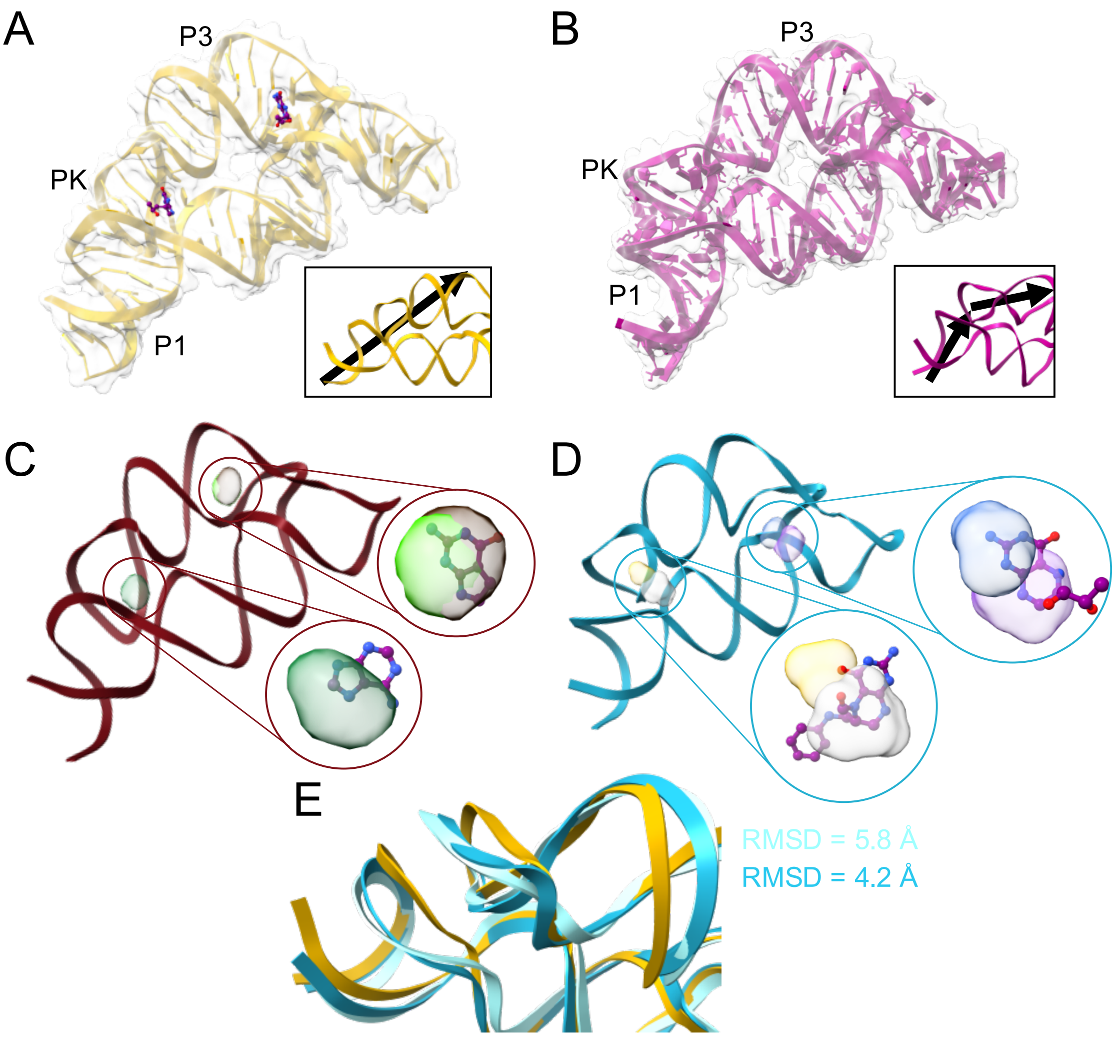

**Figure S9.** *The case of the THF riboswitch.* **AB)** Cartoon-surface representation of the THF riboswitch in its bound (**A**, PDB 4lvx[6]) and unbound (**B**, PDB 7kd1[7]) states. P1 and P3 denote the helical regions while PK denotes the pseudoknot in the molecule. The experimental ligands of the THF riboswitch are visualized in CPK style. In the inset, the relative orientation of the pseudoknot (PK) and the adjacent helical domains is highlighted by arrows. **CD)** In left panels, cartoon representation of the main conformation explored during our SHAMAN simulations started from the holo-like (**C**) and apo (**D**) states. The SHAMAPs identifying experimental binding sites in our validation set (Tab. S3) are visualized as solid surfaces with the color code defined in Tab. S7 and S8. In the insets, the SHAMAPs are overlapped with the experimental ligands: **C**) from left to right, ADE (PDB 4lw0[6]) and 7DG (PDB 4lvw[6]); **D**) from left to right, FFO (PDB 3sd3[8]) and H4B (PDB 4lvx[6]). Hydrogen atoms are visualized only when resolved in the experimental structure. **E)** Superposition of the holo (yellow ribbon), initial (cyan ribbons) and most populated (blue ribbon) structures found in the apo simulation, with focus on the pseudoknot region.

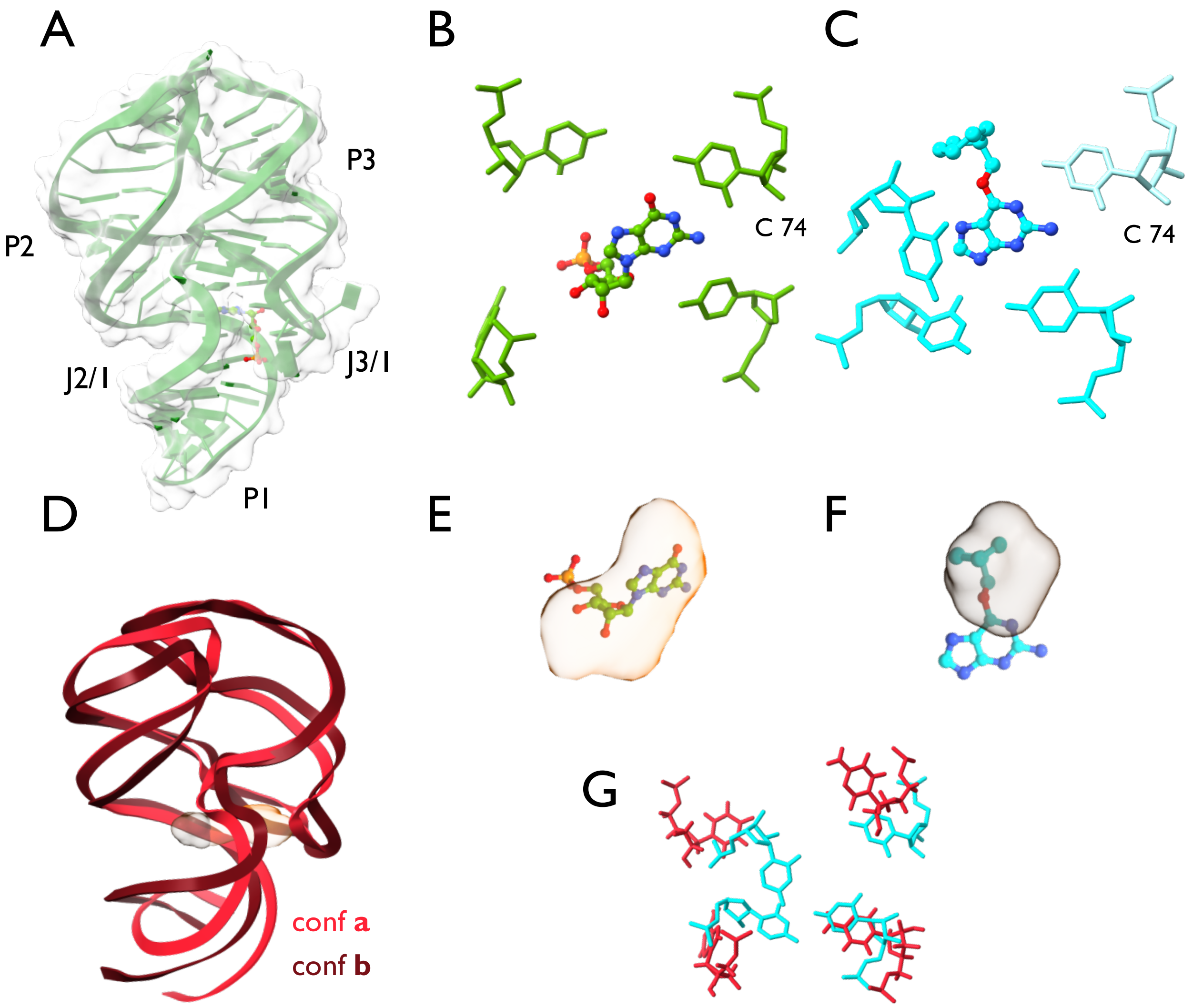

**Figure S10.** *The case of the dG riboswitch.* **A)** Cartoon-surface representation of dG riboswitch in the bound conformation, as in PDB 3slq[9]. P1-3 indicate the helical regions, while J2/1 and J3/1 the multi-way junctions. The 5GP ligand is visualized in CPK style. **BC)** The position of the key residues of interaction in the cognate (**B**, PDB 3slq with ligand 5GP) and alternative (**C**, PDB 6uc9 with ligand CMG[10]) bound states of the dG riboswitch. **D.** Cartoon representation of the two main conformations (conf **a** and **b**) explored by SHAMAN started from the holo-like state of the dG riboswitch. The SHAMAPs identifying the experimental binding site are visualized as solid surfaces with the color code defined in Tab. S7 and S8. **EF)** Superposition of the SHAMAPs with the 5GP (**E**) and CMG (**F**) ligands in the corresponding experimental structures. Hydrogen atoms are visualized only when resolved in the experimental structure. **G)** Superposition of the key interacting residues in conf **a** (red sticks) with the alternative bound structure of the dG riboswitch, after alignment of the binding regions (Materials and Methods).

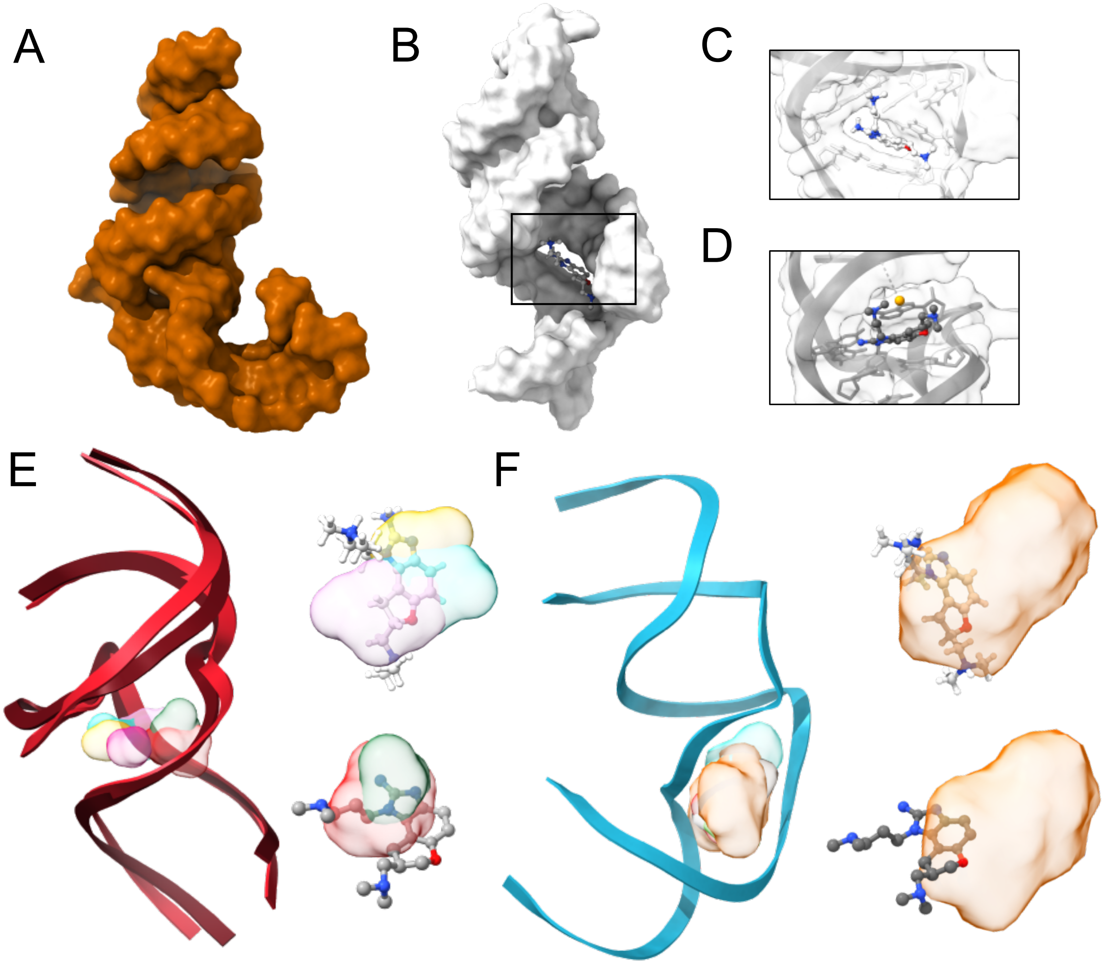

**Figure S11.** *The case of the HCV IIa IRES RNA.* **AB)** Surface representation of the HCV RNA in its apo (**A**, PDB 2nok[11]) and holo (**B**, PDB 2ku0[12]) conformations. The ISI ligand of the IAV promoter is visualized in CPK style. **CD)** Focus on the binding modes of HCV RNA bound to the ISI (PDB 2ku0[12], **C**) and SS0 (PDB 3tzr[13], **D**) ligands. Resolved magnesium ions are represented by orange spheres. **EF)** In left panels, cartoon representation of the main conformations explored during our SHAMAN simulations started from the holo-like (**E**) and apo (**F**) states of the HCV IRES IIa RNA. The SHAMAPs identifying experimental binding sites in our validation set (Tab. S4) are visualized as solid surfaces with the color code defined in Tab. S7 and S8. In the insets, the SHAMAPs are overlapped with the experimental ligands: **C**) from top to bottom, SS0 (PDB 3tzr[13]) and ISI (PDB 2ku0[12]); **D)** from top to bottom SS0 (PDB 3tzr[13]) and ISH (PDB 2ktz[12]). Hydrogen atoms are visualized only when are resolved in the experimental structure.

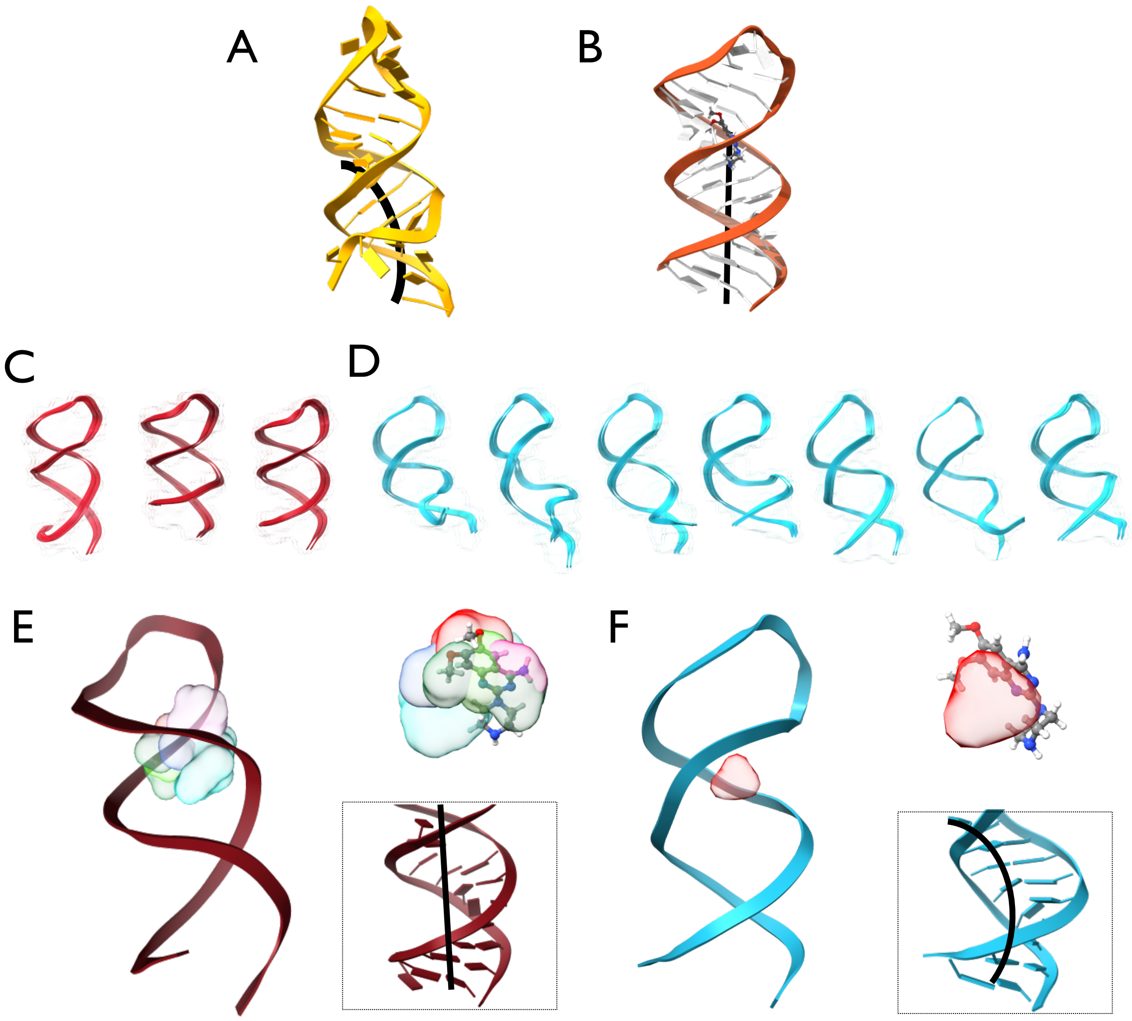

**Figure S12.** *The case of the IAV promoter.* **AB)** Cartoon representation of the HCV RNA in its apo (**A**, PDB 1mfy[14]) and bound (**B**, PDB 2lwk[15]) states. Black arrows highlight the alignment of the helical domains. The 0EC ligand of the IAV promoter is reported with CPK style. **CD)** The conformations explored during the holo-like (**C**) and apo (**D**) SHAMAN simulations of the IAV promoter. **EF)** In the left panels, cartoon representation of the conformation in which the experimental binding site was identified in the holo-like (**E**) and apo (**F**) simulations. The SHAMAPs are visualized as solid surfaces with the color code defined in Tab. S7 and S8. In the top right panel, the corresponding SHAMAPs are superimposed to the 0EC ligand. Hydrogen atoms are visualized only when resolved in the experimental structure. In the low right panel, the bending of the interhelical regions of the IAV promoter is highlighted by black arrows.

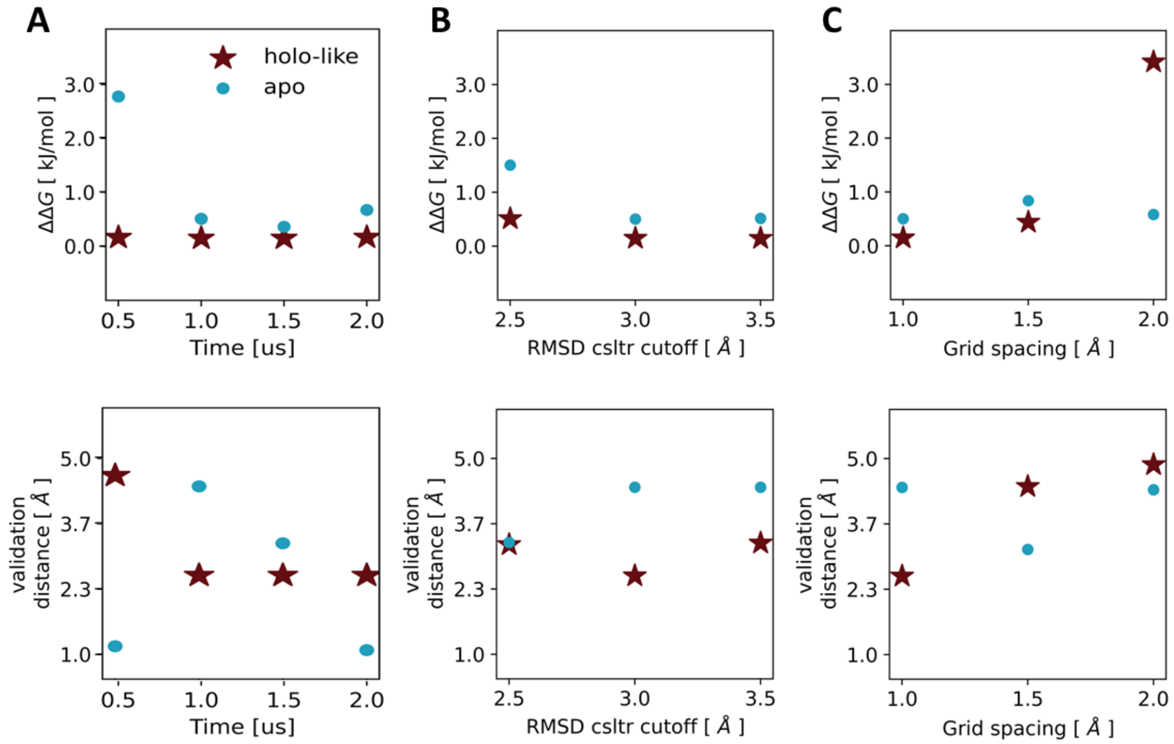

**Figure S13.** *Effect of the choice of input parameters on SHAMAN accuracy.* **A-C)** Scatter plot of ΔΔG (top panels) and validation distance (Eq. 10, bottom panels) in simulations initiated from holo-like (red) and apo (blue) conformations upon variation of: **A**) simulation time, **B**) clustering cutoff, and **C**) grid spacing in the probe binding free energy calculations. When one input parameter is varied, the other two are kept at their reference value used in the SHAMAN simulations (T = 1 µs, RMSD cutoff = 3.0 Å, grid spacing = 1.0 Å). The data reported in this figure refer to SHAMAN simulations performed on the HIV-1 TAR PDB structure 1uts.

### Supplementary analysis

##### TPP riboswitch

The thiamine pyrophosphate (TPP) riboswitch is a highly conserved riboswitch found in archaea, bacteria and eukaryotes, which directly modulates gene expression through a variety of mechanisms[16]. Upon binding to its cognate partner, the TPP, in the core region of two multi-way junctions between four helices (Fig. S8A), the TPP riboswitch assumes a stable three-dimensional structure[17]. Fragment-based screening experiments revealed the binding of several fragments in the same region in which TPP binds, but with an unexpected conformational change of residue G72, a key element in the recognition of the pyrophosphate[18] (Fig. S8B). These results indicate that alternative conformations of the TPP riboswitches may be targeted in drug design efforts, making this molecule an interesting case study for our pipeline.

We tested SHAMAN starting from a holo-like conformation of the TPP riboswitch (PDB 3d2v[19]). During this simulation, the TPP riboswitch populated a single structural cluster. The top-scored SHAMAP identified in this state (Tab. S3) corresponds to the region of the TPP binding site (left panel, Fig. S8C). The geometric accuracy in identifying the binding site resolved in the experimental structure is also impressive: the best match was obtained with ligand HPA (PDB 4nyd[5]), with a validation distance (Eq. 10) of 0.64 Å, corresponding to the best overlap of our benchmark (Tab. S3 and S4). Remarkably, our probe methyl-pyrimidine (MEPY) is perfectly overlapping with the aromatic rings of the fragment used in the aforementioned experimental screening (right panel, Fig. S8C). Importantly, a second SHAMAP with very good scoring ($\Delta\Delta G < 0.1$ kJ/mol), identified mostly by formamide (FORM) and acetate (ACEY) probes (Fig. S8D), corresponds to the region that interacts with the tail of the TPP ligand in the cognate bound state (Fig. S8B). This case study is important since, while our SHAMAN simulation captured the conformation of the TPP-bound state (red sticks in Fig. S8D), it was also able to identify the alternative binding modes by allowing for the rearrangement of the involved residues.

**THF riboswitch**

The tetrahydrofolate (THF) riboswitch, primarily found in bacteria, regulates the expression of genes involved in the synthesis and transport of the THF vitamin, which is essential for bacterial metabolism[20]. The THF riboswitch in its functional state forms two characteristic binding hotspots (Fig. S9A). Interestingly, the two pockets are both key factors of the riboswitch regulatory function and their formation and stability are interconnected[6]. In the unbound conformation of the THF riboswitch, the absence of ligands causes the unwinding of the pseudoknot PK and the misalignment of P1 and P3 helices[7] (Fig. S9B). The intrinsic structural dynamics of riboswitches and the dual-ligand binding capability of the THF riboswitch makes it an interesting case study for our pipeline.

We tested SHAMAN starting from two structures of the THF riboswitch, one in holo-like (PDB 4lvx [6]) and one in apo (PDB 7kd1[7]) conformation. In both cases SHAMAN was able to identify both binding sites among the most probable binding pockets (left panels of Fig. S9CD, Tab. S3). Interestingly, in both apo and holo-like cases the two pockets are identified within the same RNA conformation. The geometric accuracy is very high for all the identified pockets, with an average validation distance (Eq. 10) of 1.92 Å for both holo-like and apo cases (Tab. S3 and S4). The latter case is particularly remarkable since the binding helical regions in the starting apo structure are not coaxially aligned. By focusing on the P1 and P3 helices, their relative conformation explored during SHAMAN simulations (Fig. S9E, blue ribbons) resembles that present in the bound state (Fig. S9E, golden ribbons) significantly more than in the starting structure (Fig. S9E, cyan ribbons). This result exemplifies the main strength of our approach, which can capture binding pockets formed upon major conformational rearrangements.

\

**dG riboswitch**

The deoxyguanosine (dG) riboswitch is an RNA molecule found in bacteria and is involved in the regulation of metabolism by modulating their gene expression[20]. The binding site of the dG riboswitch is deeply buried in the core of a three-way junction between P1, P2 and P3 helical domains (Fig. S10A). Within this hydrophobic region, the dG riboswitch is able to recognize and bind the deoxyguanosine nucleoside *via* stable and specific base-paring interactions[20] (Fig. S10B). Several studies have revealed that the dG riboswitch is also able to undergo conformational changes and to form more solvent-exposed pockets[21]. In such pockets, other ligands may bind and, due to the absence of sugar moieties, disrupt the cognate hydrogen bonding patterns (Fig. S10C). In our validation set (Materials and Methods, Tab. S3), these two different binding modes were considered as distinct pockets. This ability of recognizing both cognate and non-cognate ligands through conformational rearrangements makes the dG riboswitch an interesting target for therapeutics approaches and an important case study for our method.

We tested SHAMAN starting from a holo-like conformation of the dG riboswitch after removal of its cognate ligand (PDB 3ski[22]). Interestingly, this riboswitch showed a relatively high flexibility, populating 5 distinct conformations in the timescale of our simulation. The two different experimental pockets were identified in two different RNA clusters, in both cases accurately characterizing the buried region of interaction (Fig. S10D). The pocket of the cognate ligand was identified by a top scored SHAMAP ($\Delta\Delta G < 0.1$ kJ/mol, Tab. S3), which is 1.6 Å away from the position of the experimental ligand and constituted solely by the BENF probe (Fig. S10E). Such result is remarkable since BENF presents similar chemical characteristics with respect to the cognate binding partners (Fig. 3). The alternative binding mode was identified by a SHAMAP with lower score ($\Delta\Delta G = 3.3$ kJ/mol), but with good geometric accuracy (3.3 Å, Fig. S10F). While our starting structure was derived from our cognate ligand bound conformation, SHAMAN simulations explored a conformation (red sticks in Fig. S10G) locally more similar to the alternative binding mode (cyan sticks in Fig. S10G). This important result demonstrates the power of our pipeline in identifying binding pockets after conformational changes of the RNA target.

**HCV IRES IIa**

The Hepatitis C Virus (HCV) Internal Ribosome Entry Site (IRES) IIa RNA is a crucial element of the HCV viral genome, as it facilitates the initiation of HCV proteins synthesis. Its functioning mechanism is peculiar: while the activity of most viral RNAs depends on the presence of translation initiation factors, HCV IRES IIa directly interacts with the host ribosomal machinery[23]. The recruitment of the ribosomal subunit to the HCV RNA is driven by its ordered folding in a L-shaped bent conformation, stabilized by divalent metal ions[11] (Fig. S11A). Stabilizing alternative conformations of the HCV RNA might therefore alter its capability to recognize the host ribosome and disrupt the viral replication, making this molecule an interesting therapeutics target. Several small molecules have been identified to bind HCV RNA[12] and to induce conformational changes that decrease the affinity to the ribosome (Fig. S11B). Among the available structures of HCV RNA bound to ligands, we identified two distinct binding sites located in the same region within the central groove of the RNA helix, but with different binding modes (Fig. S11CD, Tab. S4). The characteristic structural dynamics of the HCV RNA makes this molecule an important case study for our approach.

We tested SHAMAN starting from two conformations of the HCV IRES IIa RNA, one holo-like (PDB 3tzr[13]) and one apo (PDB 2nok[11]). In both simulations, the RNA molecule showed a higher stability compared to other viral systems studied, such as the HIV-1 TAR. This is suggested by the exploration of only two and one relevant structural clusters in the holo-like and apo simulations, respectively (left panels in Fig. EF). This stability may be attributed to the presence, in both apo and holo starting structures of HCV IRES IIa, of divalent metal ions simulated in the mother replica of SHAMAN runs (Materials and Methods). In the holo-like case, SHAMAN was able to identify both experimental binding sites (right panel, Fig. S11E) as the most probable ones ($\Delta\Delta G = 0$ and $\Delta\Delta G=0.2$ kJ/mol, respectively) and with high geometric accuracy (2.67 and 2.31 Å, respectively). In the latter case, despite the molecule populated most of the time a bent conformation close to the apo state, SHAMAN was able to detect both experimental binding sites (right panel, Fig. S11F) among the most probable ones ($\Delta\Delta G = 2.9$ and $\Delta\Delta G < 0.1$ kJ/mol, respectively), and with a slightly lower accuracy (2.84 and 3.20 Å, respectively). This case study demonstrates the ability of our approach to correctly identify the interacting hotspots independently of the starting structure of the target.

**IAV promoter**

The Influenza A Virus (IAV) promoter, positioned in the 5' untranslated region (5' UTR) of the viral genome, is a crucial player in the virus replication cycle[24]. The IAV promoter exhibits a distinctive partial duplex structure, characterized by a panhandle-like shape of the two bent helical domains (Fig. S12A). This conformation is selectively recognized by the viral polymerase, which then initiates viral transcription. In its sole available bound structure, the IAV promoter has been resolved in complex with the 0EC small molecule[15]. The latter stabilizes an alternative conformation of IAV promoter, characterized by the widening of the major RNA groove and the coaxial alignment of the angle between the helical regions (Fig. S12B). Such conformation has a lower affinity to bind the IAV polymerase enzyme and causes the inhibition of the viral activity. The challenging nature of IAV promoter and the relation between its dynamics and functions make it an important case study for our method.

We tested SHAMAN starting from two structures of the IAV promoter, one in holo-like (PDB 2lwk[15]) and one in apo (PDB 1mfy[14]) conformation. Our simulations indicated that this RNA molecule is highly flexible and populated three and seven different structural clusters in the holo-like (Fig. S12C) and apo case (Fig. S12D), respectively. The region that exhibited the highest flexibility is the lower domain of the IAV promoter: the bending of the angle between helical regions varied significantly among all the explored conformations (Fig. S12D). In the holo-like simulation, the experimental pocket was identified in the most populated cluster among the best scored SHAMAPs ($\Delta\Delta G = 0.2$ kJ/mol, left panel of Fig. S12E). Oppositely, during the apo simulation, the experimental pocket was identified in a scarcely populated cluster and with lower score ($\Delta\Delta G = 5.9$ kJ/mol, left panel Fig. S12F). The geometric accuracy was remarkable in both cases: the experimental pocket was identified with a validation distance (Eq. 10) of 1.43 and 3.64 Å, respectively (top right panel, Fig. S12EF). The lower accuracy of the apo simulation results can be explained by the structural dynamics of the helical domains. While in the holo simulation the two helical regions of the IAV promoter form a coaxial alignment characteristic of the bound state (lower right panel, Fig. S12E), in the apo simulation the interhelical angle is bent and more similar to the unbound state of IAV promoter (lower right panel, Fig. S12F). Overall, SHAMAN results for IAV promoter highlight its ability to correctly identify binding hotspots in unfavorable conditions where the target molecule is stacked in a conformation that resemble the unbound state.

#### Supplementary References

1. Panei, F. P., Torchet, R., Ménager, H., Gkeka, P. & Bonomi, M. HARIBOSS: a curated database of RNA-small molecules structures to aid rational drug design. *Bioinformatics* **38**, 4185–4193 (2022).

2. Kognole, A. A., Hazel, A. & MacKerell, A. D. SILCS-RNA: Toward a Structure-Based Drug Design Approach for Targeting RNAs with Small Molecules. *J Chem Theory Comput* **18**, 5672–5691 (2022).

3. Rizvi, N. F. *et al.* Discovery of Selective RNA-Binding Small Molecules by Affinity-Selection Mass Spectrometry. *ACS Chem Biol* **13**, 820–831 (2018).

4. Edwards, T. E. & Ferré-D'Amaré, A. R. Crystal Structures of the Thi-Box Riboswitch Bound to Thiamine Pyrophosphate Analogs Reveal Adaptive RNA-Small Molecule Recognition. *Structure* **14**, 1459–1468 (2006).

5. Warner, K. D. *et al.* Validating Fragment-Based Drug Discovery for Biological RNAs: Lead Fragments Bind and Remodel the TPP Riboswitch Specifically. *Chem Biol* **21**, 591–595 (2014).

6. Trausch, J. J. & Batey, R. T. A Disconnect between High-Affinity Binding and Efficient Regulation by Antifolates and Purines in the Tetrahydrofolate Riboswitch. *Chem Biol* **21**, 205–216 (2014).

7. Wilt, H. M., Yu, P., Tan, K., Wang, Y. X. & Stagno, J. R. Tying the knot in the tetrahydrofolate (THF) riboswitch: A molecular basis for gene regulation. *J Struct Biol* **213**, 107703 (2021).

8. Trausch, J. J., Ceres, P., Reyes, F. E. & Batey, R. T. The Structure of a Tetrahydrofolate-Sensing Riboswitch Reveals Two Ligand Binding Sites in a Single Aptamer. *Structure* **19**, 1413–1423 (2011).

9. Pikovskaya, O., Polonskaia, A., Patel, D. J. & Serganov, A. Structural principles of nucleoside selectivity in a 2′-deoxyguanosine riboswitch. *Nature Chemical Biology 2011 7:10* **7**, 748–755 (2011).

10. Matyjasik, M. M., Hall, S. D. & Batey, R. T. High Affinity Binding of N2-Modified Guanine Derivatives Significantly Disrupts the Ligand Binding Pocket of the Guanine Riboswitch. *Molecules 2020, Vol. 25, Page 2295* **25**, 2295 (2020).

11. Dibrov, S. M. *et al.* Functional Architecture of HCV IRES Domain II Stabilized by Divalent Metal Ions in the Crystal and in Solution. *Angewandte Chemie International Edition* **46**, 226–229 (2007).

12. Paulsen, R. B. *et al.* Inhibitor-induced structural change in the HCV IRES domain IIa RNA. *Proc Natl Acad Sci U S A* **107**, 7263–7268 (2010).

13. Dibrov, S. M. *et al.* Structure of a hepatitis C virus RNA domain in complex with a translation inhibitor reveals a binding mode reminiscent of riboswitches. *Proc Natl Acad Sci U S A* **109**, 5223–5228 (2012).

14. Lee, M. K. *et al.* A single-nucleotide natural variation (U4 to C4) in an influenza A virus promoter exhibits a large structural change: implications for differential viral RNA synthesis by RNA-dependent RNA polymerase. *Nucleic Acids Res* **31**, 1216–1223 (2003).

15. Lee, M. K. *et al.* A novel small-molecule binds to the influenza A virus RNA promoter and inhibits viral replication. *Chemical Communications* **50**, 368–370 (2013).

16. Subki, A., Ho, C. L., Nabihan Ismail, N. F., Zainal Abidin, A. A. & Balia Yusof, Z. N. Identification and characterisation of thiamine pyrophosphate (TPP) riboswitch in Elaeis guineensis. *PLoS One* **15**, e0235431 (2020).

17. Antunes, D., Jorge, N. A. N., Garcia de Souza Costa, M., Passetti, F. & Caffarena, E. R. Unraveling RNA dynamical behavior of TPP riboswitches: a comparison between Escherichia coli and Arabidopsis thaliana. *Scientific Reports 2019 9:1* **9**, 1–13 (2019).

18. Warner, K. D. *et al.* Validating Fragment-Based Drug Discovery for Biological RNAs: Lead Fragments Bind and Remodel the TPP Riboswitch Specifically. *Chem Biol* **21**, 591–595 (2014).

19. Thore, S., Frick, C. & Ban, N. Structural basis of thiamine pyrophosphate analogues binding to the eukaryotic riboswitch. *J Am Chem Soc* **130**, 8116–8117 (2008).

20. Matyjasik, M. M. & Batey, R. T. Structural basis for 2′-deoxyguanosine recognition by the 2′-dG-II class of riboswitches. *Nucleic Acids Res* **47**, 10931–10941 (2019).

21. Matyjasik, M. M., Hall, S. D. & Batey, R. T. High Affinity Binding of N2-Modified Guanine Derivatives Significantly Disrupts the Ligand Binding Pocket of the Guanine Riboswitch. *Molecules 2020, Vol. 25, Page 2295* **25**, 2295 (2020).

22. Pikovskaya, O., Polonskaia, A., Patel, D. J. & Serganov, A. Structural principles of nucleoside selectivity in a 2′-deoxyguanosine riboswitch. *Nature Chemical Biology 2011 7:10* **7**, 748–755 (2011).

23. Lukavsky, P. J. Structure and function of HCV IRES domains. *Virus Res* **139**, 166–171 (2009).

24. Noble, E. *et al.* Biophysical Analysis of Influenza A Virus RNA Promoter at Physiological Temperatures. *Journal of Biological Chemistry* **286**, 22965–22970 (2011).
